# Influenza A M2 Inhibitor Binding Understood through Mechanisms of Excess Proton Stabilization and Channel Dynamics

**DOI:** 10.1101/2020.06.19.162248

**Authors:** Laura C. Watkins, William F. DeGrado, Gregory A. Voth

**Affiliations:** Department of Chemistry, Chicago Center for Theoretical Chemistry, Institute for Biophysical Dynamics and James Franck Institute, The University of Chicago, Chicago, Illinois 60637, United States; Department of Pharmaceutical Chemistry, University of California, San Francisco, San Francisco, California, 94158, United States

## Abstract

Prevalent resistance to inhibitors that target the influenza A M2 proton channel has necessitated a continued drug design effort, supported by a sustained study of the mechanism of channel function and inhibition. Recent high-resolution X-ray crystal structures present the first opportunity to see how the adamantyl-amine class of inhibitors bind to M2 and disrupt and interact with the channel’s water network, providing insight into the critical properties that enable their effective inhibition in wildtype M2. In this work, we test the hypothesis that these drugs act primarily as mechanism-based inhibitors by comparing hydrated excess proton stabilization during proton transport in M2 with the interactions revealed in the crystal structures, using the Multiscale Reactive Molecular Dynamics (MS-RMD) methodology. MS-RMD, unlike classical molecular dynamics, models the hydrated proton (hydronium-like cation) as a dynamic excess charge defect and allows bonds to break and form, capturing the intricate interactions between the hydrated excess proton, protein atoms, and water. Through this, we show that the ammonium group of the inhibitors is effectively positioned to take advantage of the channel’s natural ability to stabilize an excess protonic charge and is thus acting as a hydronium-mimic. Additionally, we show that the channel is especially stable in the drug binding region, highlighting the importance of this property for binding the adamantane group. Finally, we characterize an additional hinge point near Val27, which dynamically responds to charge and inhibitor binding. Altogether, this work further illuminates a dynamic understanding of the mechanism of drug inhibition in M2, grounded in the fundamental properties that enable the channel to transport and stabilize excess protons, with critical implications for future drug design efforts.

**TOC Graphic:** 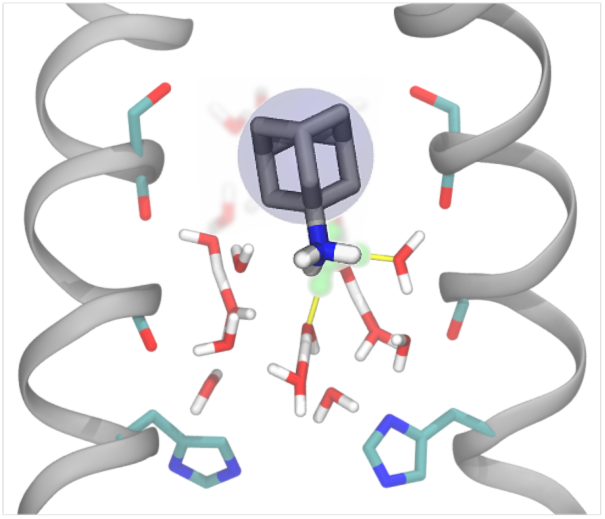

## INTRODUCTION

Proton transport (PT) across cellular membranes is a critical component of many biomolecular systems, necessary, for example, to maintain pH gradients,^1, 2^ to drive ATP synthesis,^3^ and to facilitate the co- or anti-transport of other small molecules.^4-6^ Because of their essential role in such systems, channels and transporters with PT functionality are often targets for drug design to inhibit or control PT—in the case of viruses and bacteria, to slow or prevent infection, but there are myriad other disease applications.^7-9^ Drug design is notoriously challenging, as both thermodynamic and kinetic factors must be considered but are difficult to predict and control, and its success depends on high quality structures, an understanding of structural dynamics, and a knowledge of the protein’s function and its mechanism. Thus, beyond elucidating mechanisms of PT in order to understand how a specific channel or transporter works, studying the detailed interactions that facilitate PT can provide valuable insight to help guide drug design efforts.

The influenza virus kills up to 650,000 people each year,^10^ and the impact of the recent global coronavirus pandemic^11^ emphasizes how critical it is to maintain our focus on understanding and treating viral infections. The influenza A virus matrix 2 (M2) proton channel is a homo-tetrameric protein responsible for the acidification of the viral interior, a critical step in the influenza infection process.^12-14^ It is the target of two of the three currently available oral antivirals, amantadine and rimantadine.^15, 16^ While these are effective at blocking PT in wildtype M2, drug-resistant mutants have become the predominate strains, rendering these drugs ineffective and thus requiring a continued drug design effort^17^ informed by a deeper understanding of the PT and drug inhibition mechanisms. Additionally, M2 is considered an archetype for the viroporin family, a class of viral channels considered ideal drug targets.^18^ The SARS-CoV-2 virus responsible for the COVID-19 pandemic contains two viroporins, protein E and 3.^19-21^ Thus, viroporins are a critical class of proteins to study as potential therapeutic targets.

M2 is located in the viral capsid and is acid-activated: as the pH of the endosome encapsulating the virus is lowered, the M2 channel becomes activated and facilitates unidirectional proton flow to the viral interior, allowing the virus to escape the endosome and infect the cell. The key residue that controls activation is His37,^22-24^ which can bind one additional proton and take on a +1 charge. One histidine from each helix forms the His37 tetrad, which can collectively hold a +0 to +4 excess charge, dependent on pH. The channel becomes activated and the C-terminal portion opens (adopting the Inward_open_ conformation) upon reaching the +3 state, and PT occurs as the channel cycles through a transporter-like mechanism.^25-32^

Amantadine and rimantadine belong to the adamantyl-amine class of inhibitors, binding in the upper-middle portion of the channel. These drugs were the predecessors of many related adamantane-based compounds featuring a relatively rigid, apolar group and an attached charged group.^33-39^ Recently, Thomaston et al. published several high-resolution X-ray crystal structures of M2 with amantadine, rimantadine, and a novel spiro-adamantyl amine bound.^40^ These structures provided the first opportunity to see the specific interactions that facilitate stable inhibitor binding and the disruption of the hydrogen-bonded water network otherwise present. Along with an earlier qualitative MD simulation study that guided the design of the spiro-adamantyl amine inhibitors,^41^ the crystallographic analysis provided potential insights into the mechanism of inhibition, suggesting that the backbone carbonyls of pore-lining residues act as physiochemical chameleons, able to engage in both hydrophobic and hydrophilic interactions, and that the drug is tilted off the channel’s axis and interacts with waters in the Ala30 layer. Taken together, it is hypothesized that amantadine acts as a mechanism-based inhibitor, with the ammonium group functioning as a hydronium mimic. Computational studies to date have primarily focused on the means of entry into the channel and location of binding,^42-46^ but have not deduced specific interactions between the drug, channel, and channel water involved in binding as they relate specifically to similar interactions seen in the PT mechanism.

Proton transport is an inherently quantum mechanical process, as the hydrated proton structure (hydronium-like) exists in a complex hydrogen-bonded network that rearranges dynamically as bonds break and form according to the Grotthuss shuttling mechanism.^47-49^ Thus, classical molecular dynamics (MD) with fixed bonding topology cannot be used to study PT; moreover, *ab initio* methods are not efficient enough to reach the many nanosecond timescales necessary to obtain sufficient sampling in biomolecular systems that may have important degrees of freedom several orders of magnitude slower than proton shuttling. Multiscale Reactive Molecular Dynamics (MS-RMD)^50-53^ (and Multistate Empirical Valence Bond, MS-EVB, before it) was developed to efficiently and accurately capture the solvation and delocalization of an excess proton in water, such that the quantum-chemical nature of the hydrated proton can be studied in the context of membrane proteins over the long timescales needed for accurate simulation of such systems. MS-RMD has been successfully applied in several protein systems to predict and explain mechanisms of PT.^31, 32, 54-61^

In previous work,^31, 32^ quantum mechanics/molecular mechanics (QM/MM) and MS-RMD was used to calculate potentials of mean force (PMFs, i.e., free energy profiles) of PT through the M2 channel in the +0-3 states, providing critical insight into the pH-dependent activation behavior and the role of the His37 tetrad in PT. Most recently, we further analyzed the MS-RMD simulations to explore the detailed interactions between the hydrated excess proton and the channel and found that the proton dynamically, as a function of its position, alters several properties of the protein and pore waters, including the hydrogen-bonding network and the protein structure.^61^ This latter work illustrates how MS-RMD can be used successfully to investigate explicit, dynamic interactions between a hydrated proton and its immediate environment, as well as its indirect effects on other parts of the system. Here, we employ a similar approach as in this previous work to focus specifically on properties related to drug binding and how the position of the bound drug relates to the overall PT mechanism. Through this analysis, we examine the hypothesis that the adamantyl-amine drugs act as mechanism-based inhibitors. Our results indicate that the ammonium group of amantadine and rimantadine are aptly positioned to take advantage of the channel’s ability to stabilize an excess proton. Additionally, by examining conformational fluctuations, we show that the drug binding pocket is an especially stable and symmetrical portion of the channel, conducive to binding a roughly spherical drug, and we reveal an additional minor hinge point towards the top of the channel which may be a relevant feature for future drug design efforts.

## METHODS

Simulations for calculating properties as the proton moves through the top of the channel were run as follows. Starting structures were taken from previous simulations, which were initiated from a crystal structure of the transmembrane portion of the M2 channel (this construct is referred to as M2TM) resolved at room-temperature and high pH (PDB: 4QKL^62^) embedded in a 1-palmitoyl-2-oleoyl-sn-glycero-3-phosphocholine (POPC) bilayer solvated with water. M2TM is the minimum construct necessary to retain proton conduction similar to full-length M2,^63^ and it has been shown that the presence of amphipathic helices, included in the full-length M2 protein, do not significantly influence the PT mechanism.^32^ The collective variable (CV) defined for umbrella sampling (US) is the z-coordinate of the vector between the excess proton center of excess charge (CEC, see below) and the center of mass of the four Gly34 alpha carbons, as in our previous work, such that the CV has negative values at the top (N-terminal end) of the protein and progresses to positive values at the bottom (C-terminal end). The excess proton CEC is defined as:^64^

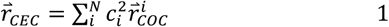

where 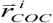 is the center-of-charge of the diabatic MS-RMD state, and 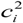 is the amplitude of that state. The sum is over all *N* states. MS-RMD simulations were run with the excess proton at every 0.5 Å along the CV coordinate between −18.0 and 1.0, generating 39 windows. To ensure that the proton remained in the channel, a cylindrical restraint was added at 8 Å with a force constant of 10 kcal/mol·Å^2^ using the open-source, community-developed PLUMED library.^65, 66^ The MS-EVB version 3.2 parameters^67^ were used to describe the hydrated excess proton. After a 250 ps MS-RMD equilibration, the replica exchange umbrella sampling^68^ technique was used to facilitate convergence. Production simulations were run for ∼2-4 ns with frames saved every 1 ps.

For calculating hydrogen bond residence times, longer independent trajectories were run with the CEC restrained in 5 different positions using US as described above. Each trajectory was run for 1.75-2 ps with frames saved every 10fs.

Simulation frames were binned by excess proton CEC value for subsequent analyses, which were performed in Python^69^ using the SciPy,^70^ NumPy,^71^ and pandas^72^ libraries. For hydrogen bond analysis, values were averaged over the four helices. Hydrogen bonds were defined by the following criteria: the donor-acceptor distance must be less than 3.5Å, and the donor-hydrogen-acceptor angle must be greater than 150°. Several hydrogen bond definitions were tested and did not affect the conclusions (not shown). For calculating residence times, a hydrogen bond was considered in place as long as the particular water molecule remained the closest water to the protein atom and the hydrogen bond criteria were met.

Images of molecular structures were rendered in Visual Molecular Dynamics (VMD),^73^ while other figures were generated using Matplotlib.^74^

## RESULTS AND DISCUSSION

If the adamantyl-amine drugs are acting as mechanism-based inhibitors as hypothesized, we would expect to see specific aspects of the PT mechanism taken advantage of or replicated by the drug upon drug binding. To test this, we performed MS-RMD simulations of M2 in the +0 His37 charge state with an explicit excess proton to evaluate the hydrogen-bonding networks, pore shape, and protein fluctuations throughout PT that relate to drug binding. By focusing on PT in the +0 state, we are studying the process of proton entry and diffusion to His37 in the first key step of channel activation, paralleling inhibitor entry into the channel. We additionally expect this to be a prevalent charge state in drug-bound structures due to the lowered His37 pKas.^75^ Replica exchange umbrella sampling was used to obtain sufficient sampling of proton positions throughout the top portion of the channel, with windows from CEC_Z_ = −18.0 to 1.0 Å where the coordinate origin is defined as center of mass of the Gly34 alpha carbons. The channel is aligned along the z-axis for all subsequent analyses.

To understand how properties of PT may provide insight into drug binding, we primarily examine variations dependent on proton position. We compared the values of each property when the proton is at the drugs’ ammonium group positions versus other parts of the channel to determine if the drugs could be taking advantage of the channel’s natural ability to stabilize a proton. This idea is highlighted in Figure 1, which shows both the drug-bound crystal structure and a snapshot of a hydrated excess proton in the channel from our simulations. We refer to the drugs’ ammonium nitrogen position along the z-axis in the crystal structure as AmmN_Z_. This value is −1.7 and −1.5 Å for the the Inward_closed_ amantadine and rimantadine bound structures, respectively (averaged over the two tetramers in each crystal structure).

**Figure 1.**
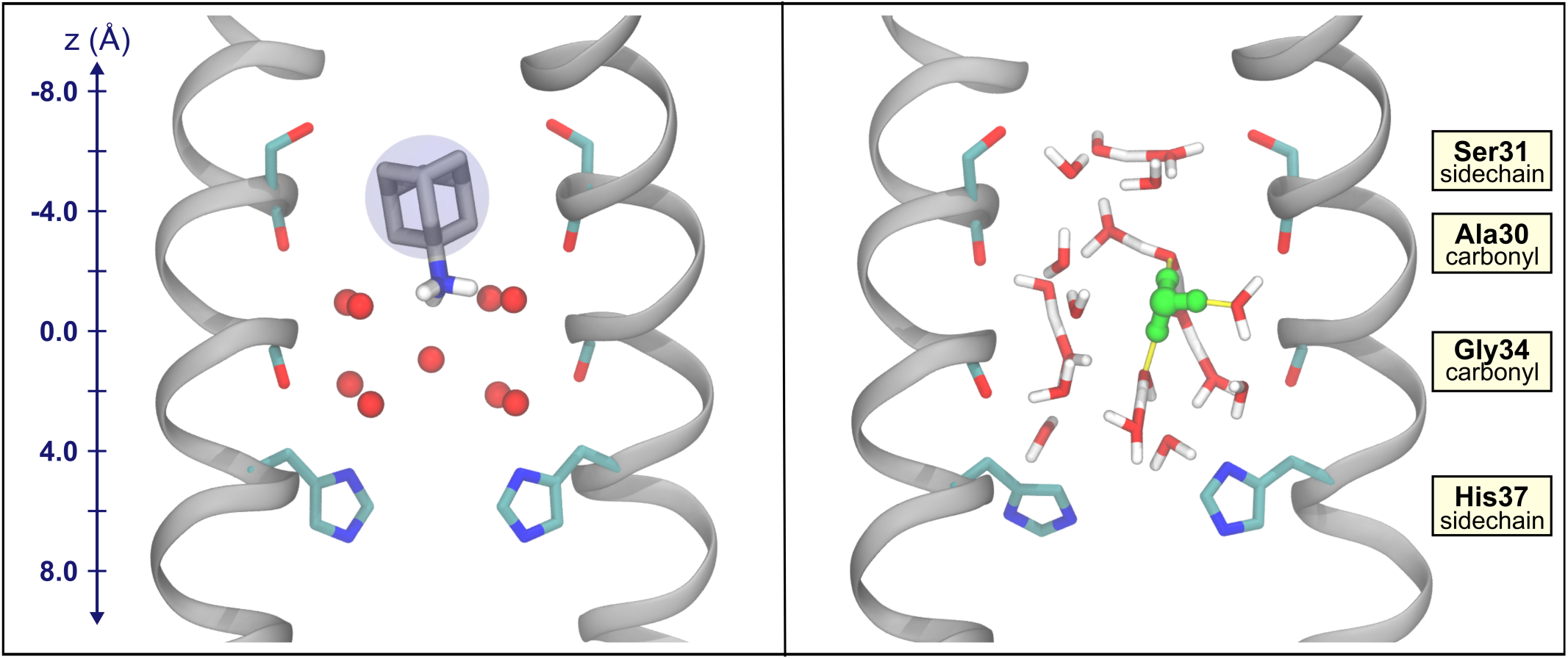
Comparison of M2 with amantadine and the hydrated excess proton. Left, x-ray crystal structure of amantadine-bound M2, PDB 6BKK. Water oxygens are shown in red. A blue circle around the adamantane group is depicted to reflect the spherical nature of this group. Hydrogens on the ammonium group of amantadine were added. Right, a snapshot from an MS-RMD trajectory with the most hydronium-like water indicated in green. In both, two opposing chains of M2 are shown in silver. The Ser31, His37 sidechains and the Gly34, Ala30 backbone carbonyls are shown. The z-coordinate for the system is included on the left, where z = 0 Å is defined as the center-of-mass of the Gly34 alpha-carbons.

### Flexible Hydrogen Bonds Stabilize the Excess Proton near AmmN_Z_

It has been shown in our previous work that hydrogen bonds within the channel, including those between water and protein atoms, help facilitate proton transport by altering their direction and frequency of interaction as the proton moves through the channel. Here, we focus specifically on water interactions that may help account for excess charge stabilization near AmmN_Z_. In Figure 2, we calculate the occupancy of three different hydrogen bonds between protein atoms and water as a function of the excess proton position in the channel. While the Ala30 hydrogen bond occupancy is consistent as the excess proton enters and moves through the top of the channel, as it approaches the Ala30 carbonyls, the occupancy decreases ∼20%. This dip indicates the Ala30 hydrogen-bonded waters can flexibly reduce their interaction with the protein as a result of an excess charge in their vicinity. Additionally, this dip is centered at −2.5 Å, near AmmN_Z_. At this point, the role of the waters near Ala30 carbonyls in hydrating the proton is maximized. This supports the hypothesis that amantadine and rimantadine are mechanism-based inhibitors and take advantage of the channel’s natural ability to stabilize a hydrated excess proton in order to stabilize the drug’s ammonium group.

**Figure 2.**
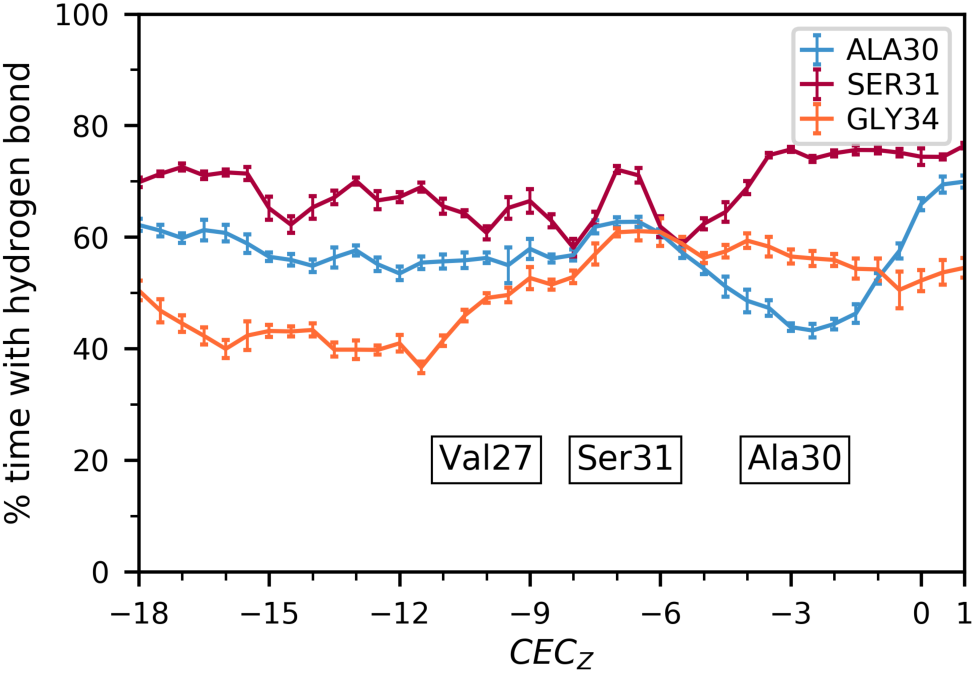
Hydrogen bond occupancy averages and error as a function of hydrated excess proton position between water and the backbone carbonyls of Ala30, Gly34, and the sidechain hydroxyl group of Ser31. Approximate average positions of Val27 sidechains, Ser31 sidechains, and Ala30 carbonyls are indicated.

The hydrogen bond occupancy of waters with the Gly34 carbonyls increases once the excess proton passes through the Val27 gate and remains fairly consistent across proton positions thereafter, exhibiting little dependence on the hydrated proton position once it is in the channel. The Ser31 sidechain water occupancies are shown for comparison, which do not show a noticeable trend based on proton position. Thus, this change in interactions is not a universal effect throughout the channel, but the Ala30 waters seem to be uniquely flexible in this manner. These differences are consistent with drug design studies —while compounds such as spiro-adamantyl amine have been able to displace the water in the Ala30 layer, no designed inhibitors have displaced the water around Gly34.

To further understand how the dynamics of the hydrogen-bond network may show how these drugs benefit from the channel’s inherent excess-charge stabilization used in proton transport, we examined the average residence times of hydrogen bonds between water and several important protein atoms. To do this, independent trajectories were run for five different excess proton positions, including two trajectories with the proton completely outside the channel (CEC_Z_ = −24.0, 24.0 Å) and three when the proton is near AmmN_Z_. These results are shown in Figure 3. The Ala30 water residence times slightly increase when the excess proton is near AmmN_Z_, the Ser31 water residence times do not show any significant difference between the proton outside the channel and at AmmN_Z_, and those of Gly34 waters decrease at AmmN_Z_. The waters hydrogen bonded to His37 imidazole nitrogens show the greatest change in residence times and are shown to highlight the ability of this method to describe such differences. We note that classical MD does not wholly capture charge transfer in hydrogen bonds, resulting in an overall weaker interaction. Thus, while these simulations provide valuable insight into these interactions and their trends, we expect that they are stronger in the real system and any differences will be more prominent.

**Figure 3.**
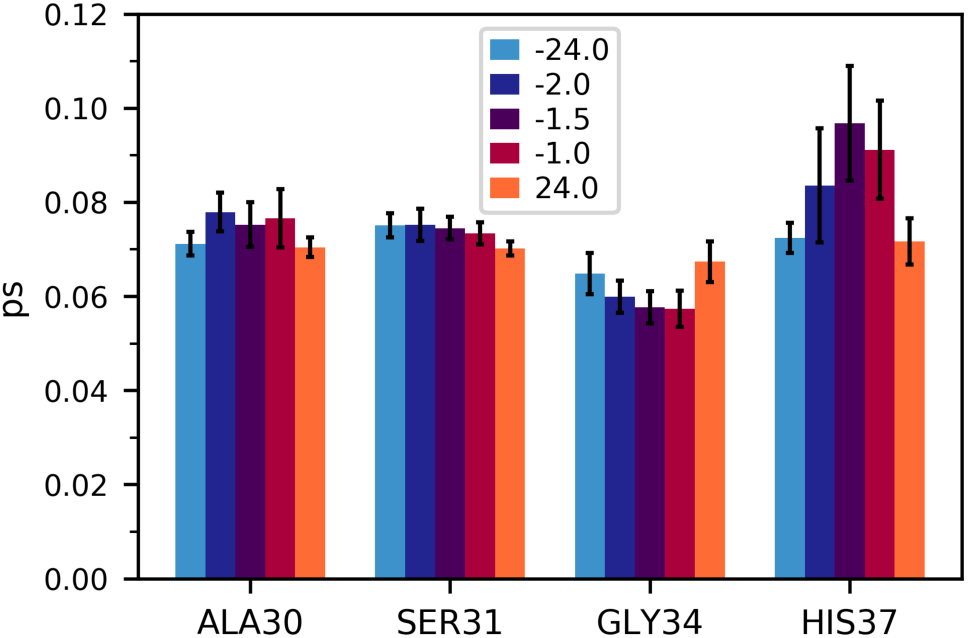
Average hydrogen bond residence times, calculated for the backbone carbonyls if Ala30, Ser31, and Gly34, and the unprotonated nitrogen of the His37 imidazole sidechain. The quantities are averaged over the interactions of all four helices, and error bars were calculated using block averaging. Each bar represents the value calculated from one trajectory with the proton restrained at the labeled Z coordinate.

With the above results for Ala30, this may indicate that several waters remain tightly hydrogen bonded to the Ala30 backbone carbonyls, while one or more are bonded less frequently. While the Gly34 hydrogen bonds do not form less frequently (as indicated in Figure 2) with an excess charge in this region, they do exhibit greater dynamics and flexibility. This change indicates an increase in water dynamics when the excess proton is near, which could help stabilize and solvate the excess charge in the Ala30 water layer.

Taken together, these results further support the hypothesis that amantadine and rimantadine act as mechanism-based inhibitors: the channel acts as a scaffold to facilitate PT by harboring flexible protein-water interactions that can adapt and respond to a positive excess charge, with specific ability to stabilize an excess proton near AmmN_Z_ that the charged drugs can take advantage of.

### Drug Tilt Positions Ammonium Group in Highest CEC Density

One prominent characteristic of the amantadine and rimantadine bound structures is the drug’s tilt within the pore. This tilted conformation is also seen in solid-state NMR studies.^76^ Given the drug’s three-fold symmetry in a four-fold symmetric channel, the ammonium group cannot form hydrogen bonds with all 4 waters hydrogen-bonded to Ala30, leading in part to this tilt. Based on our previous work examining the proton’s path through the channel, we used a similar analysis to examine the density of CEC positions when the excess proton is near the ammonium position in drug-bound structures. Figure 4A shows the difference in CEC density when the proton is near AmmN_Z_ compared with the average over all proton positions through the top portion of the channel. This 2-dimensional histogram of CEC positions in the XY-plane is calculated for CEC_Z_ = −1.7±0.2 Å, minus the average over all normalized histograms for CEC_Z_ positions [-18.0, 1.0] Å binned every 0.2 Å. Interestingly, in this portion of the channel the excess proton prefers to be near the edge of the pore, unlike the predominate preference for the center of the pore throughout the rest of the channel, as indicated by the positive values around the edge and negative values in the center. Figure 4B shows the radial density of the CEC in this same region of the channel. Possible positions of amantadine’s ammonium group nitrogens were calculated based on the drug’s position and tilt in the crystal structure, and their radii are included as dashed lines (these positions were also used to generate the image in Figure 1). These hydrogens can extend to a radius ∼1.5 Å in this static crystal structure, which indicates that the slightly off-centered ammonium group directly positions 1-2 of its hydrogens in the region of the CEC’s highest density. The CEC’s propensity for the edge of the pore indicates that the drug’s tilt in the channel may not only be a necessary component of its binding, but also a thermodynamic advantage. This tilt further allows the drug to act as a hydronium mimic, as the hydrogens of the ammonium group are in the favorable positions of the solvated excess proton.

**Figure 4.**
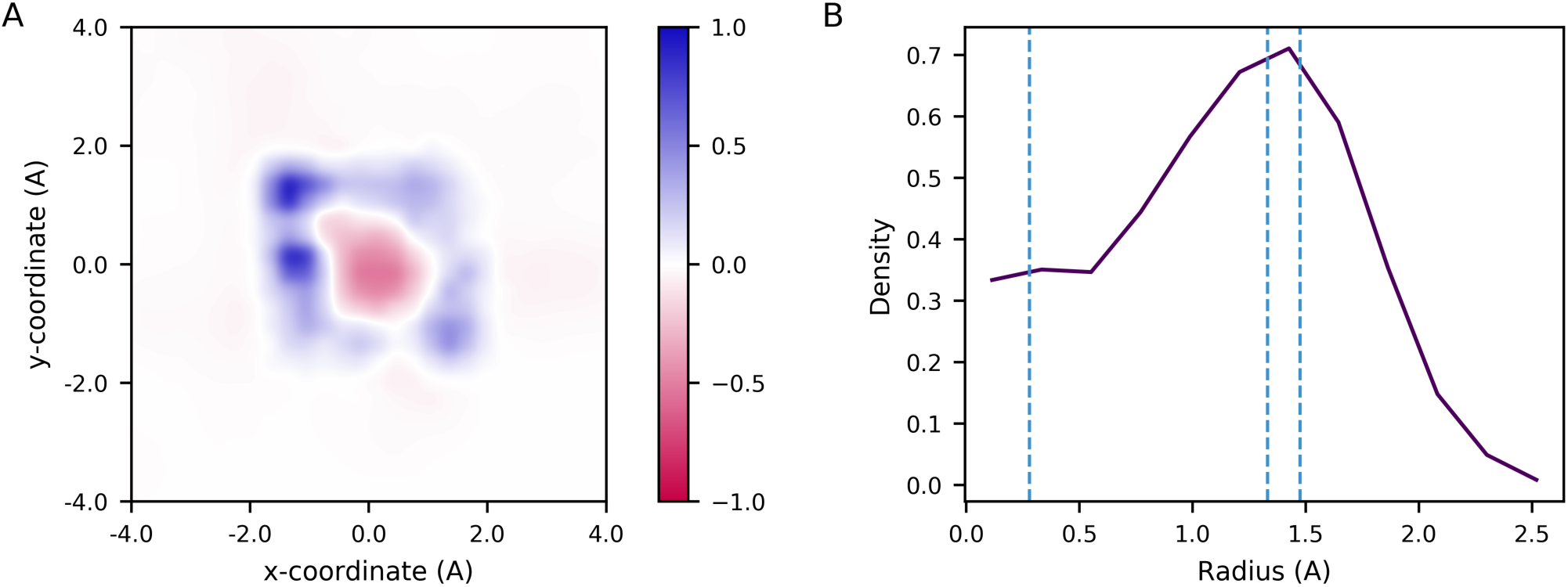
A. The difference in hydrated excess CEC density in the xy-plane when the CEC is located within CEC_Z_=-1.7±0.2 Å compared with the average CEC density over all CEC positions in range z=[-18.0,1.0]. B. The average radial density of the proton when CEC_Z_ =- 1.7±0.2 Å. One possible set of positions of amantadine’s ammonium group hydrogens calculated from the drug-bound crystal structure are indicated by blue dashed lines.

The analysis of hydrogen bonding changes and proton densities indicates how the ammonium group is a functional addition to the adamantane scaffold, as the charged group is positioned in a region where the channel is especially adept to stabilize an excess charge. This stabilization relies on flexible water structures and hydrogen-bond interactions that can undergo minor changes to accommodate the proton, suggesting that the adamantyl-amine inhibitors are acting as hydronium mimics — they take advantage of these inherent features to help solvate the charged ammonium group. The identification of other regions of the channel with increased ability to stabilizing an excess charge, such as areas of increased proton density or significantly flexible water interactions, could help provide new targets for drugs to act as hydronium-mimics.

### Pore Shape and Stability Near Ser31 are Ideal for Adamantane Binding

Another hypothesis about the adamantyl-amine class of inhibitors is that adamantane is effectively spherical and can freely rotate within the channel, but has no rotatable bonds, which minimizes the entropy lost upon binding. This rapid rotation can be seen on the NMR time scale^76^ and is consistent with the recent Thomaston et al. crystallographic studies, in which the motion was indirectly inferred. Nevertheless, its significance depends on the dynamic nature of the channel — if the protein exhibits great structural fluctuations in the region where the drug binds, then drug binding may induce changes that greatly decrease the entropy and this hypothesis would not fully explain the drugs’ efficacy. To better understand how the channel’s natural dynamics may lend itself to favorable drug binding, we examined the pore shape throughout our trajectories. As an estimate of the asymmetry of the channel, we calculated the eccentricity, which is essentially a measure of how “circular” a given oval is. The eccentricity is defined as:

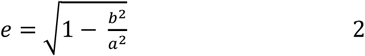

where *a* and *b* are the semi-major and semi-minor axes, respectively, which we approximate by the distance between alpha-carbons on opposing helices. A schematic of this is shown in SI Figure 1. Eccentricity can have values between 0 and 1, with 0 indicating a circle and 1 indicating a parabola.

These results are shown in Figure 5. Eccentricity maximum, minimum, and RMSD values are calculated in SI Table 1. The pore-lining residues in the bottom half of the channel, Gly34, His37, and Trp41, all show a greater degree of asymmetry and a wider range of eccentricity values, dependent on the excess proton position, than the pore-lining residues in the top part of the channel. Interestingly, proton entry at CEC_z_ = −17 Å has a pronounced effect on the channel near Trp41, greater than that when the proton nears the center of the channel. Ser31, however, has overall the smallest average eccentricity and the lowest minimum value than the other pore-lining residues during PT in this portion of the channel. Additionally, Ser31 and Val27 have smaller proton position dependent changes in eccentricity than the pore-lining residues in the bottom half of the channel. This result indicates that the Ser31 region is the most symmetrical and stable in the channel.

**Figure 5.**
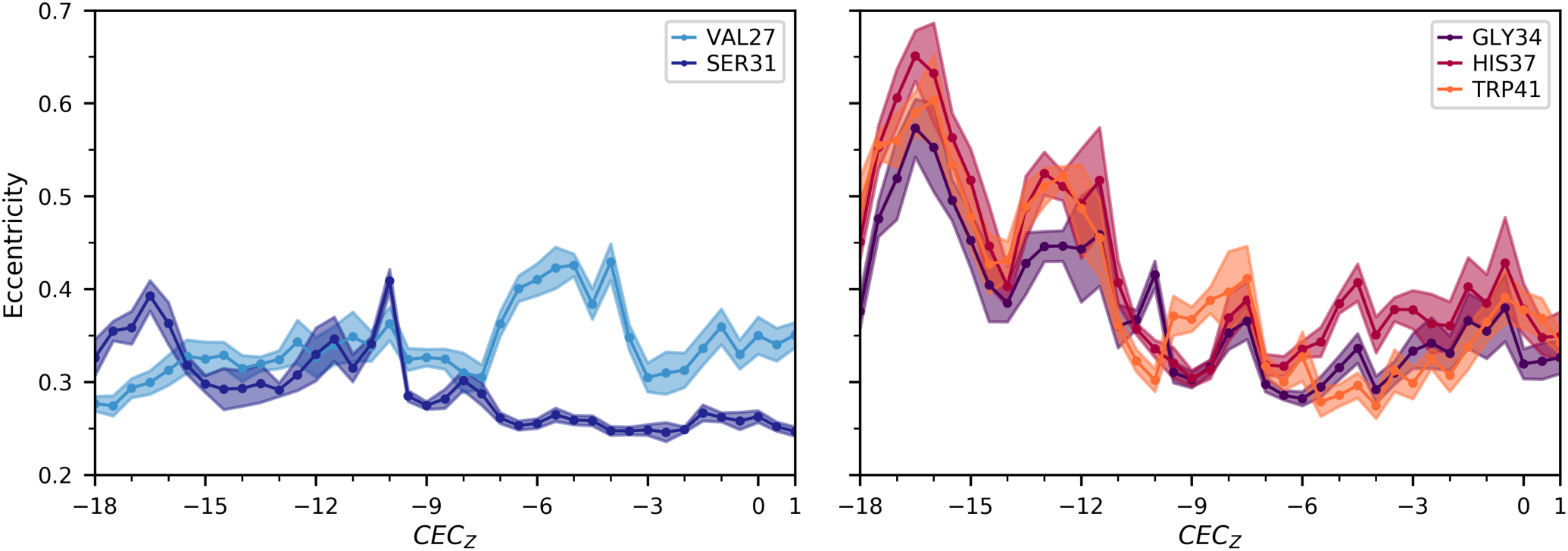
Average and standard deviation of the pore eccentricity estimated by alpha-carbon distances of the pore-lining residues as a function of the hydrated excess proton CEC position.

While analyzing the alpha-carbon distances and eccentricity, we also examined the correlation between these alpha-carbons distances on opposing helices, shown in Figure 6, to gain further insight into protein motion and conformational fluctuations on the nanosecond timescale. These motions captured here are equilibrium fluctuations in the +0, Inward_closed_ state, not necessarily motions driving the transition between Inward_open_ and Inward_closed_. The calculated correlations indicate that the channel’s equilibrium structural fluctuations are dominated by alternating inward-outward motions of opposing helices. At each pore-lining residue, the distances are negatively correlated—that is, when helices A and C move farther apart, helices B and D move closer together, and vice-versa.

**Figure 6.**
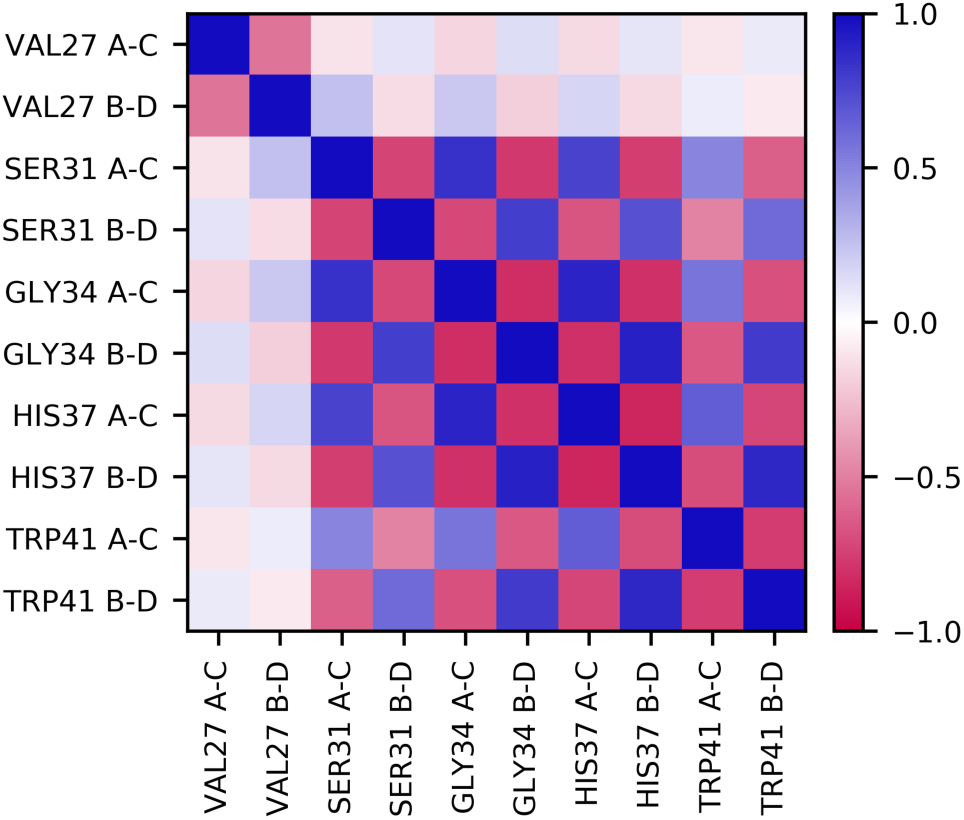
Pearson correlation coefficients of the distances between alpha-carbons on opposing helices, for all pore-lining residues, when excess proton CEC_Z_=-18.0Å. Each row and column correspond to a specific residue and distance, as labeled, where ‘A-C’ is the distance between the alpha-carbons of helices and 1 and 3, and ‘B-D’ is that of helices 2 and 4. Only those values with a p-value < 0.05 are shown, any other values are set to 0.0.

Gly34 is known to be the hinge point whose kinking controls the large structural change between inward-open and inward-closed conformations, which may falsely lead to the conclusion that the conformational fluctuations at equilibrium above and below Gly34 are decorrelated, with a stable core centered at Gly34. Interestingly, however, the motions at Gly34 are strongly and similarly correlated with the motions at both Ser31 and His37. This correlation indicates that in this fixed charge state, the Gly34 kink is relatively rigid. Instead, there is a noticeable lack of correlation between Val27 and the other pore-lining residues, suggesting that there is a secondary, minor “hinge” between Val27 and Ser31 that decorrelate the inward-outward motions between the helices above and below this point. This natural hinge observed near Val27 furthers our understanding of Val27 acting as a secondary gate that opens to allow proton and water entry into the channel.^77, 78^ In our simulations, this valve can readily hydrate, particularly in the presence of a nearby excess proton. Moreover, it is frequently closed, which may make passage of a hydrated sodium or chloride ion more difficult. This aspect of the Val27 gate and its relevance for PT and proton selectivity is likely an important feature of the M2 channel and could be further explored in future work.

Given that the adamantane group of the drug is centered in the Ser31 tetrad plane, we hypothesize that these facets of the channel’s dynamics are critical for fully explaining the drugs’ favorable binding. Because of the more circular shape of the pore at the Ser31 tetrad, the spherical adamantane group can fit snugly under the hydrophobic Val27 cleft and block PT. Additionally, the relative stability of the pore in the region of drug binding helps explain why drug binding is thermodynamically favorable. Because the channel exhibits smaller structural fluctuations here than in other regions of the channel, the adamantane based drugs are able to bind with minimal loss of entropy as the channel does not need to lose flexibility to create a stable drug-binding interface. This has important implications for designing drugs that use scaffolds different from the adamantane group,^37, 79-84^ or that interact with drug-resistant mutants such as S31N. While drug binding to more flexible regions of the channel is possible, only modest changes in potency are often observed despite large changes in the size of the drugs. This is likely because of the need to counter the greater loss of entropy resultant from structural changes and reduced fluctuations.

## CONCLUSIONS

Altogether, we have shown how the adamantyl-amine inhibitors of M2 are suited to exploit various inherent features of the M2 channel that naturally facilitate proton transport, further supporting the claim that they function as mechanism-based inhibitors. The ability of hydrogen bond interactions to flexibly respond to the hydrated excess proton CEC, measured in both hydrogen bond occupancy and residence times, indicate how the channel is suited to stabilizing an excess charge near AmmN_Z_. Thus, the ammonium group of these inhibitors can act as a hydronium-mimic by binding in this region. We also analyzed the pore shape throughout the channel by calculating the eccentricity of the pore based on alpha-carbon distances. The results from these calculations indicate that the drug binding pocket is an especially stable and symmetrical portion of the channel, conducive to binding a roughly spherical drug. Finally, by examining the correlations between these distances, we found an additional minor hinge point towards the top of the channel which may be a relevant feature for future drug design efforts.

Understanding these features as they relate to drug binding gives further insight into the specific interactions that stabilize drug binding, and could help inform drug design efforts by highlighting other aspects of the M2 channel and proton transport mechanism that a drug could take advantage of. Through this understanding, we hope future drug-design efforts can take advantage of this approach to methodically create new inhibitors for the more prevalent mutant strains. With the recent publication of high resolution influenza B M2 (BM2) structures,^85^ we hope similar studies to elucidate the detailed PT mechanism in BM2 will be conducted to guide drug design in this functionally similar protein.

This work also shows how similar analyses to understand the details of explicit proton transport mechanisms (not those inferred by water structures alone) could be used in other systems, and extended to ion transporters such as the SARS-CoV-2 viroporins, to help inform mechanism-based inhibitor design. Elucidating the inherent features of drug-targetable proton transporters, such as flexible water and hydrogen bonding interactions, preferred proton positions, dynamic pore shapes, and structural fluctuations, can help guide the design of drug scaffolds and added substituents. The MS-RMD simulation methodology utilized in this work has also made these studies possible both for M2 and other important drug targets.

## ASSOCIATED CONTENT

### Supporting Information

A table of average, maximum, minimum, and RMSD for eccentricity is included in the SI. This material is available free of charge via the Internet at http://pubs.acs.org.

## AUTHOR INFORMATION

### Notes

The authors declare no competing financial interest.

## ACKNOWLEDGMENT

The personnel in this research were supported by the National Institute of General Medical Sciences (NIGMS) of the National Institutes of Health (NIH Grant R01 GM053148). The authors acknowledge the University of Chicago Research Computing Center and the U.S. Department of Defense High Performance Computing Modernization Program for providing computing resources. LCW received funding from Department of Energy (DOE) Computational Science Graduate Fellowship under grant DE-FG02-97ER25308.

## Notes

### Competing Interest Statement

The authors have declared no competing interest.

