## Supplemental File for "Influenza A M2 Inhibitor Binding Understood through Mechanisms of Excess Proton Stabilization and Channel Dynamics"

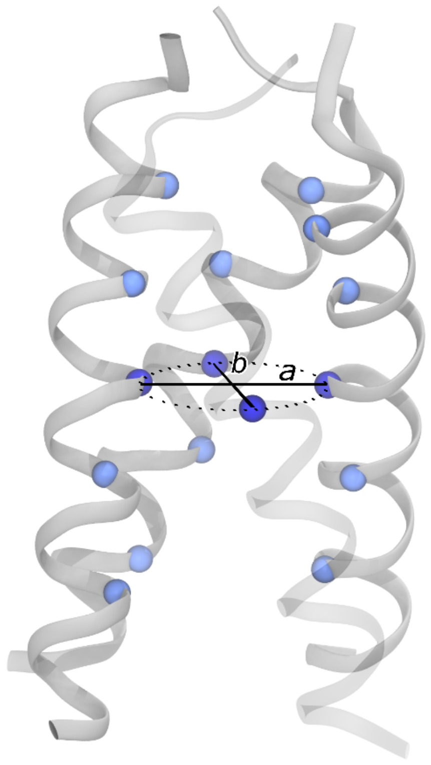


**SI Figure 1**. A schematic of eccentricity calculations, with pore-lining alpha-carbons shown as blue spheres. Dark blue spheres are Gly34 alpha-carbons. The major and minor axes of the oval are defined by the distances between alpha-carbons on opposing helices, indicated as lines *a* and *b*.

**SI Table 1**. Eccentricity values of the pore at each of the pore-lining residues, calculated using alpha-carbon positions as described in the text. Eccentricity was calculated for proton positions in the top [1]half of the channel, $CEC_z$ = [-18.0,-1.0] Å.

| Residue | Average *e* over all $CEC_z$ | Max value of *e* and position | Min value of *e* and position | Difference between max and min values | RMSD |
|---|---|---|---|---|---|
| **Val27** | 0.34 | 0.43<br>$CEC_z$ = -4.0 Å | 0.27<br>$CEC_z$ = -17.5 Å | 0.16 | 0.13 |
| **Ser31** | 0.29 | 0.41<br>$CEC_z$ = -10.0 Å | 0.25<br>$CEC_z$ = -2.5 Å | 0.16 | 0.13 |
| **Gly34** | 0.38 | 0.57<br>$CEC_z$ = -16.5 Å | 0.28<br>$CEC_z$ = -6.0 Å | 0.29 | 0.17 |
| **His37** | 0.42 | 0.65<br>$CEC_z$ = -16.5 Å | 0.30<br>$CEC_z$ = -9.0 Å | 0.35 | 0.18 |
| **Trp41** | 0.40 | 0.60<br>$CEC_z$ = -16.0 Å | 0.27<br>$CEC_z$ = -4.0 Å | 0.33 | 0.17 |

*
